# Water stabilizes an alternate turn conformation in horse heart myoglobin

**DOI:** 10.1101/2023.01.15.524139

**Authors:** Alex Bronstein, Ailie Marx

## Abstract

Comparison of myoglobin structures reveals that protein isolated from horse heart consistently adopts an alternate turn conformation in comparison to its homologues. Analysis of hundreds of high-resolution structures discounts crystallization conditions or the surrounding amino acid protein environment as explaining this difference, that is also not captured by the AlphaFold prediction. Rather, a water molecule is identified as stabilizing the conformation in the horse heart structure, which immediately reverts to the whale conformation in molecular dynamics simulations excluding that structural water.

## Introduction

Sperm whale myoglobin was the first, high resolution, protein structure ever solved by X-ray diffraction analysis (1). Today this small, globular, single-domain protein is used as a model in exploring protein folding (2). Myoglobin is represented in the Protein Data Bank (PDB) by hundreds of structures. The large number of independent experimental realizations of this protein structure provides a unique opportunity to identify features intrinsic to the protein and not the result of specific experimental artifacts. Here we observe that horse heart myoglobin consistently adopts a unique loop conformation and show, using molecular dynamics, that a structural water is necessary and sufficient for maintaining this conformation. This unique conformation highlights a structural feature which state of the art structure prediction methods such as AlphaFold remain blind to.

## Results

The hundreds of myoglobin structures available in the PDB share a very high degree of structural similarity across crystallization conditions, species, and mutations (Fig. 1A). None-the-less, variations exist, ranging from the major global conformational changes in domain swapped heterodimeric myoglobin to the local and specific adoption of different turn conformations in the interhelical region between helices G and H (Fig. 1B and 1C). The latter was noted originally in the first high resolution structural comparison between sperm whale myoglobin (*Phy*_myoglobin) and horse heart myoglobin (*Eq*_myoglobin) (3) and was attributed to an altered bonding network resulting from minor primary sequence differences. Many structures later, we suggest that sequence variations cannot clearly account for this difference. Evans and Bayer (3) noted that the H12N substitution creates an N12–K16 interaction in *Eq*_myoglobin at the expense of the K16–D122 salt bridge in *Phy*_myoglobin, and also that the D27–R118 pair in *Eq*_myoglobin form a salt bridge not found between the equivalent E27– K118 pair in *Phy*_myoglobin. However, pig myoglobin (*Sus*_myoglobin), also characterized by tens of high-resolution structures, has the same turn conformation as *Phy*_myoglobin despite having the H12N and D27–R118 sequence features of *Eq*_myoglobin (Fig. 1D). Unique to *Eq*_myoglobin is Q9 (L9 in *Phy*_myoglobin, *Sus*_myoglobin and all homologues crystallized to date) that forms a polar interaction with D126 of helix H. However, we discount this feature as directly accounting for the altered conformation since there are *Eq*_myoglobin adopting the *Phy*_myoglobin conformation, described below, and without any alteration to or around the Q9 either in sequence or space.

**Fig. 1.**
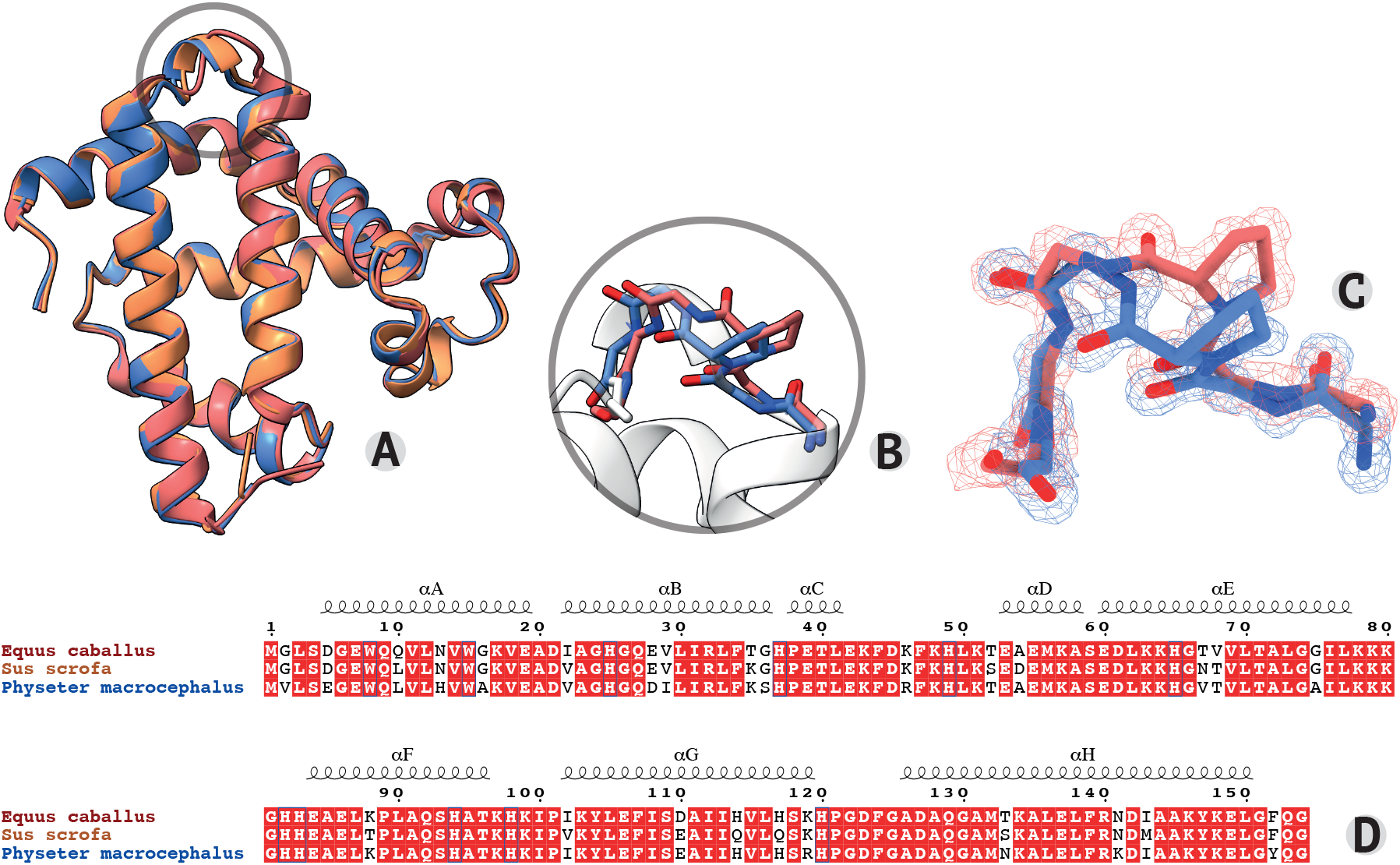
Horse heart myoglobin adopts an alternate turn conformation which is not explainable at the sequence level. *Eq*_myoglobin (1GJN, maroon), *Phy*_myoglobin (1A6K, blue) and *Sus*_myoglobin (1MWC, orange) structure alignment (A), highlighting the altered *Eq*_myoglobin turn conformation between helices G and H (B), a region with well-defined electron density(C). High sequence identity is demonstrated in alignment (D).

To track the source of the highly reproducible, alternate loop conformations we assessed all myoglobin structures. Table 1 shows that most species adopt the type I turn conformation, the clear exception being *Eq*_Myoglobin for which the type II turn is the preferred conformation. Analysis of these PDB entries verifies that crystallization conditions cannot readily explain the altered turn conformations; *Eq*_myoglobin and *Phy*_myoglobin are both represented by numerous structures crystallized in space groups P 2_1_ 2_1_ 2_1_ and P 1 2_1_ 1, and crystallized from conditions of both high salt concentration (usually ammonium sulphate) and various molecular weight polyethylene glycol (PEG) solutions.

**Table 1.**
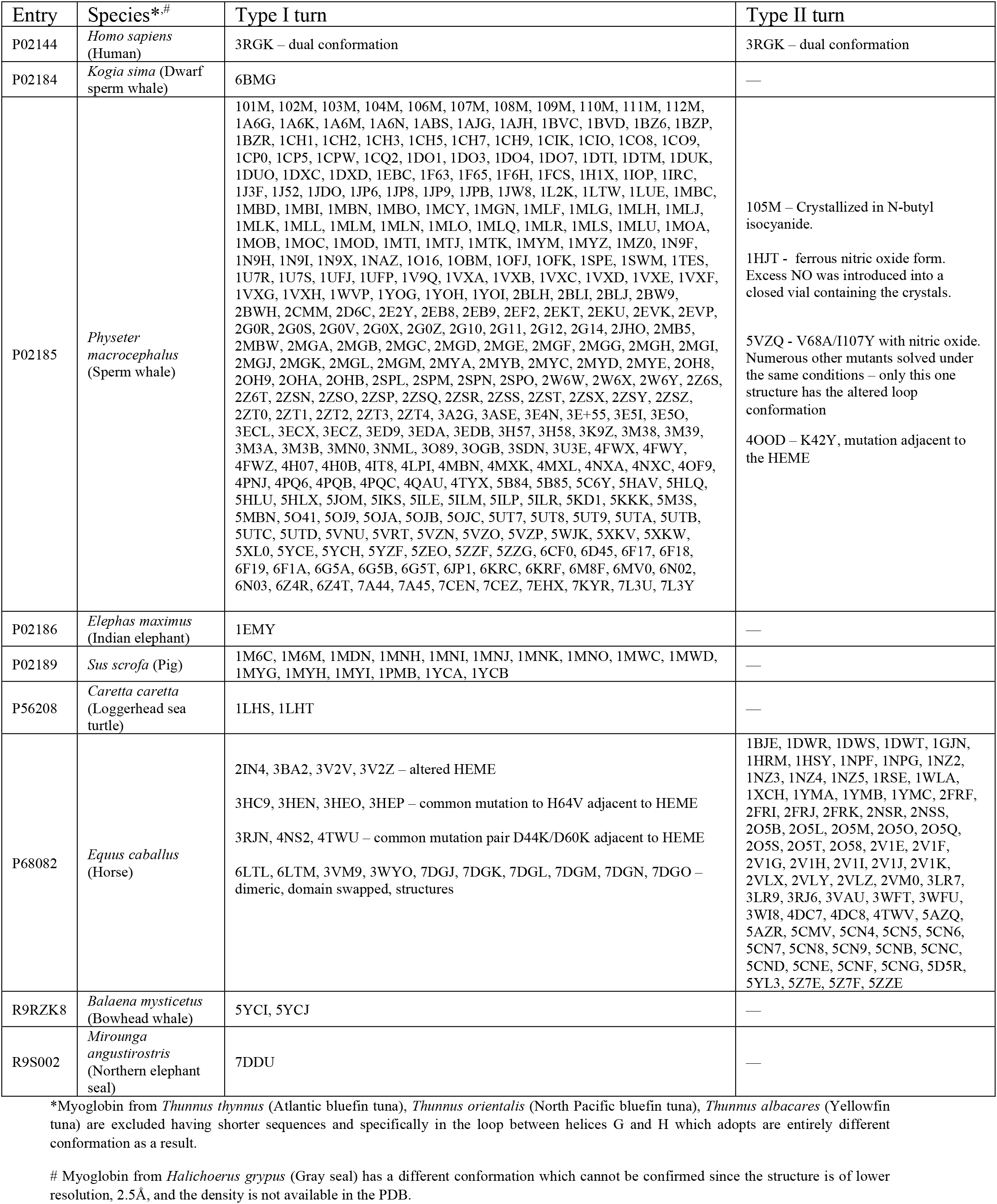
Categorization of the turn conformation adopted by myoglobin structures.

The identifying features of *Eq*_Myoglobin structures adopting the *Phy*_myoglobin are annotated in Table 1 and include (i) those having a wild type sequence but binding an altered heme cofactor (e.g. 3BA2, chlorin-substituted *Eq*_Myoglobin), (ii) those having certain mutations adjacent to the heme binding site, and (iii) heterodimeric, domain-swapped structures. The association with heme binding is curious given that the loop in question is located on the opposite end of the protein, at least 25Å away, and in a loop not observed to have an altered conformation in the apo protein (4). It was demonstrated decades ago that heme binding occurs cotranslationally (5), and also that myoglobin can bind heme in different orientations having different rate constants (6); more recently, molecular dynamics simulations showed that heme binding modulates the myoglobin folding pathway, increasing myoglobin stability and folding cooperativity (7). Studies into the folding pathways of domain-swapped variants indicate differences between the native monomer and domain-swapped protein (8). Together these observations could suggest that the variant loop conformations in *Eq*_myoglobin and *Phy*_myoglobin are products of subtly different folding pathways.

To assess if these alternate conformations are intrinsically stable within the folded protein, we first ran molecular dynamics (MD) simulations on the myoglobin protein chain alone. In these simulations the initial *Eq*_myoglobin conformation quickly (10-20ns) adjusted to and remained stable in the *Phy*_myoglobin conformation (refer to 1us simulations in Fig. 2A). Further analysis of the structures revealed a ubiquitously present water molecule, or electron density supporting water (e.g., 2VLY has a hydrogen peroxide molecule modelled in place of water, and 5AZQ has clear electron density in the 2Fo-Fc map despite no water molecule is modelled), bound in a network of three hydrogen bonds within the *Eq*_myoglobin turn conformation (Fig. 2B). MD simulations maintaining this structural water, demonstrated that its presence is sufficient to hold in place the alternate *Eq*_myoglobin loop conformation (Fig. 2A). This water molecule is not present in *Phy*_myoglobin structures or *Eq*_myoglobin having the *Phy*_myoglobin conformation, despite (spatial) conservation of the coordinating residues. Further supporting the critical role of this water in defining the local structure between helices G and H in *Eq*_myoglobin is AlphaFold2 structure prediction, which fails to account for sequence-extrinsic structural elements like water and predicts the *Eq*_myoglobin sequence in the *Phy*_myoglobin turn conformation (Fig. 2C).

**Fig. 2.**
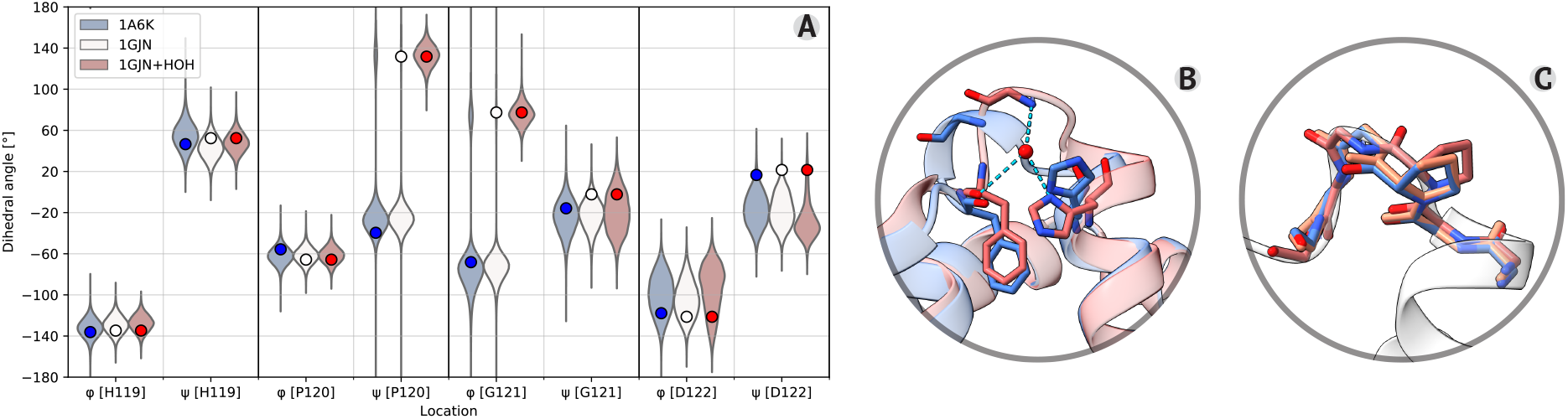
The horse heart turn conformation is associated with the presence of a structural water molecule. A display of dihedral angle distributions from 1us molecular dynamics simulations (violins) and initial values in PDB structures (circles) show that *Eq*_myoglobin (white) adopts the *Phy*_myoglobin (blue) turn conformation unless a structural water is maintained with the *Eq*_myoglobin (red) (A). This structural water forms three hydrogen bonds in the the *Eq*_myoglobin structure but not the *Phy*_myoglobin structure despite the presence of the same amino acid network (B). AlphaFold predicts *Phy*_myoglobin conformation from *Eq*_myoglobin sequence (grey cartoon, aligned to *Eq*_myoglobin 1GJN (red) and *Phy*_myoglobin 1A6K (blue)) (C).

## Discussion

We conclude that the distinct conformation adopted by the protein chain bridging helices G and H in *Phy*_myoglobin and *Eq*_myoglobin structures is consistent across a large ensemble of structures and cannot be readily explained by the sequence alone, the local environment of the protein chain, crystallization conditions or ambiguity in the structural models. When and how a water molecule, critical for maintaining the *Eq*_myoglobin turn conformation, becomes incorporated into the structure and how this is directed by mutations far removed from that location in sequence and space should be a matter for further investigations. The species specificity of this phenomenon, the observation that mutations affecting folding rate can alter the conformational preference and especially since turns are thought to be involved in the early stages of protein folding (9) may implicate a protein folding mechanism. Our analysis suggests the continued usefulness of myoglobin as a “model” for protein folding, perhaps in the exploration of co-translational folding.

## Author contributions

AB and AM contributed together to all aspects of this work. AM ran the molecular dynamics simulations.

## Declaration of interests

The authors declare no competing interests.

## Materials and methods

### Sequence alignment

Sequences were aligned using Clustalw (10) and displayed using the ESPript 3.0 server (11).

### Structure analysis

All structures associated with Uniprot (12) entries for the *MB* gene were aligned and analysed using The PyMOL Molecular Graphics System, Version 2.0 Schrödinger, LLC (13). Figures were prepared using ChimeraX (14).

### Molecular dynamics simulations

Simulations were run on apo-myoglobin using models 1GJN and 1A6K with all water molecules removed and additionally on 1GJN with water molecule 2112 held under position restraint with harmonic force constant of 100000 kJ/mol^-1^ nm^-2^. All simulations were conducted using GROMACS software, version 2020.2 (15). The Amber 99sb-ildnp force field (16) was applied to normal amino acids and ions, and the TIP3P model (17) was applied to water molecules. After the energy minimizations and heating to 300K, the system was equilibrated under NVT (constant volume and constant temperature) and NPT (constant pressure and constant temperature) conditions. Production runs were performed under NPT conditions, with a time step of 2 fs. The temperature and pressure were maintained at 300 K and 1 bar. Simulations were sampled every 20 ps for each trajectory and the distributions of each dihedral angle, for each model were displayed on a violin plot. Two simulations with different initial velocities were conducted for each system to ensure reproducibility.

## Notes

### Competing Interest Statement

The authors have declared no competing interest.

